# The Opiliones Tree of Life: shedding light on harvestmen relationships through transcriptomics

**DOI:** 10.1101/077594

**Authors:** Rosa Fernándeza, Prashant Sharma, Ana L.M. Tourinho, Gonzalo Giribet

## Abstract

Opiliones are iconic arachnids with a Paleozoic origin and a diversity that reflects ancient biogeographical patterns dating back at least to the times of Pangea. Due to interest in harvestman diversity, evolution and biogeography, their relationships have been thoroughly studied using morphology and PCR-based Sanger approaches to systematics. More recently, two studies utilized transcriptomics-based phylogenomics to explore their basal relationships and diversification, but sampling was limiting for understanding deep evolutionary patterns, as they lacked good taxon representation at the family level. Here we analyze a set of the 14 existing transcriptomes with 40 additional ones generated for this study, representing ca. 80% of the extant familial diversity in Opiliones. Our phylogenetic analyses, including a set of data matrices with different gene occupancy and evolutionary rates, and using a multitude of methods correcting for a diversity of factors affecting phylogenomic data matrices, provide a robust and stable Opiliones tree of life, where most families are precisely placed. Our dating analyses also using alternative calibration points, methods, and analytical parameters provide well-resolved old divergences, consistent with ancient regionalization in Pangea in some groups, and Pangean vicariance in others. The integration of state-of-the-art molecular techniques and analyses, together with the broadest taxonomic sampling to date presented in a phylogenomic study of harvestmen, provide new insights into harvestmen interrelationships, as well as a general overview of the general biogeographic patterns of this ancient arthropod group.

## Introduction

Opiliones are a remarkable group of arachnids (Figs 1–2), with a fossil record dating to the Early Devonian having diversified in its main lineages by the Carboniferous (Dunlop et al., 2003; Dunlop, 2007; Garwood et al., 2014), showing ancient vicariant patterns that accord with their modern distribution (Boyer et al., 2007; Giribet and Kury, 2007; Clouse and Giribet, 2010; Giribet et al., 2012a; Giribet and Sharma, 2015). They show fascinating reproductive behaviors, including paternal and biparental care (Machado, 2007; Machado and Macías-Ordóñez, 2007; Requena et al., 2012; Buzatto et al., 2014).

**Figure 1.**
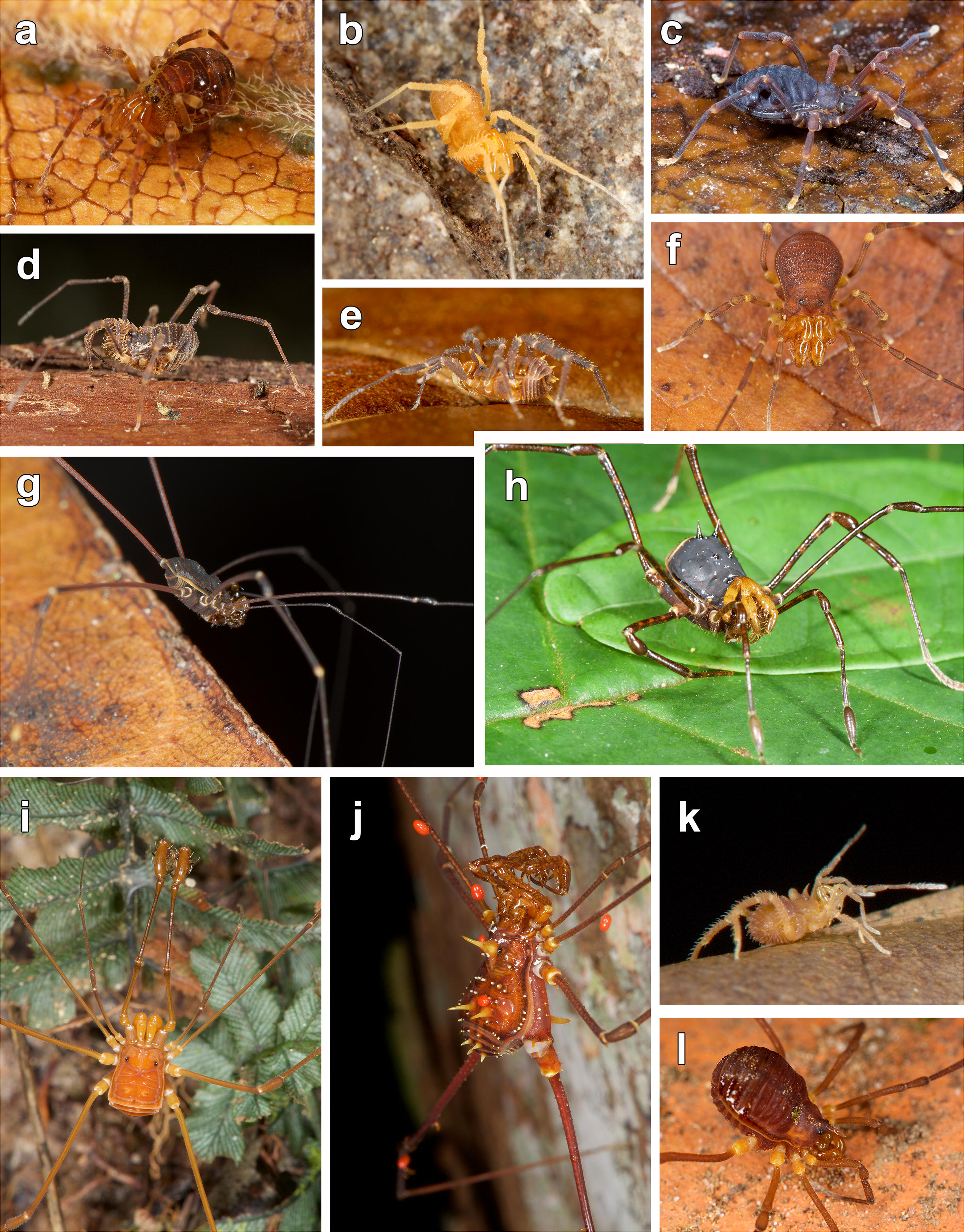
Live habitus of selected specimens: a. Aoraki longitarsa from Aoraki/Mount Cook NP, South Island, New Zealand, 20.i.2014 (MCZ 137238); b. Parasiro coiffaiti from Queralbs, Barcelona, Catalonia, Spain, 15.vii.2014 (MCZ 43732); c. Caddo pepperella from Petersham, Massachusetts, USA, 24.vi.2014 (MCZ 40441); d. Forsteropsalis pureora from Waikeremoana, North Island, New Zealand, 12.i.2014; e. Odiellus troguloides from Queralbs, Barcelona, Catalonia, Spain, 15.vii.2014 (MCZ 43657); f. Homalenotus remyi from Queralbs, Barcelona, Catalonia, Spain, 15.vii.2014 (MCZ 43715); g. Acropsopilio neozealandiae from Putai Ngahere Reserve, North Island, New Zealand, 14.i.2016 (MCZ 30457); h. Ischyropsalis nodifera from Sodupe, Bilbo, Euskadi, Spain, photographed 13.viii.2014 (MCZ 45501); i. Nemastomella dubia from Queralbs, Barcelona, Catalonia, Spain, 15.vii.2014 (MCZ 43619)

**Figure 2.**
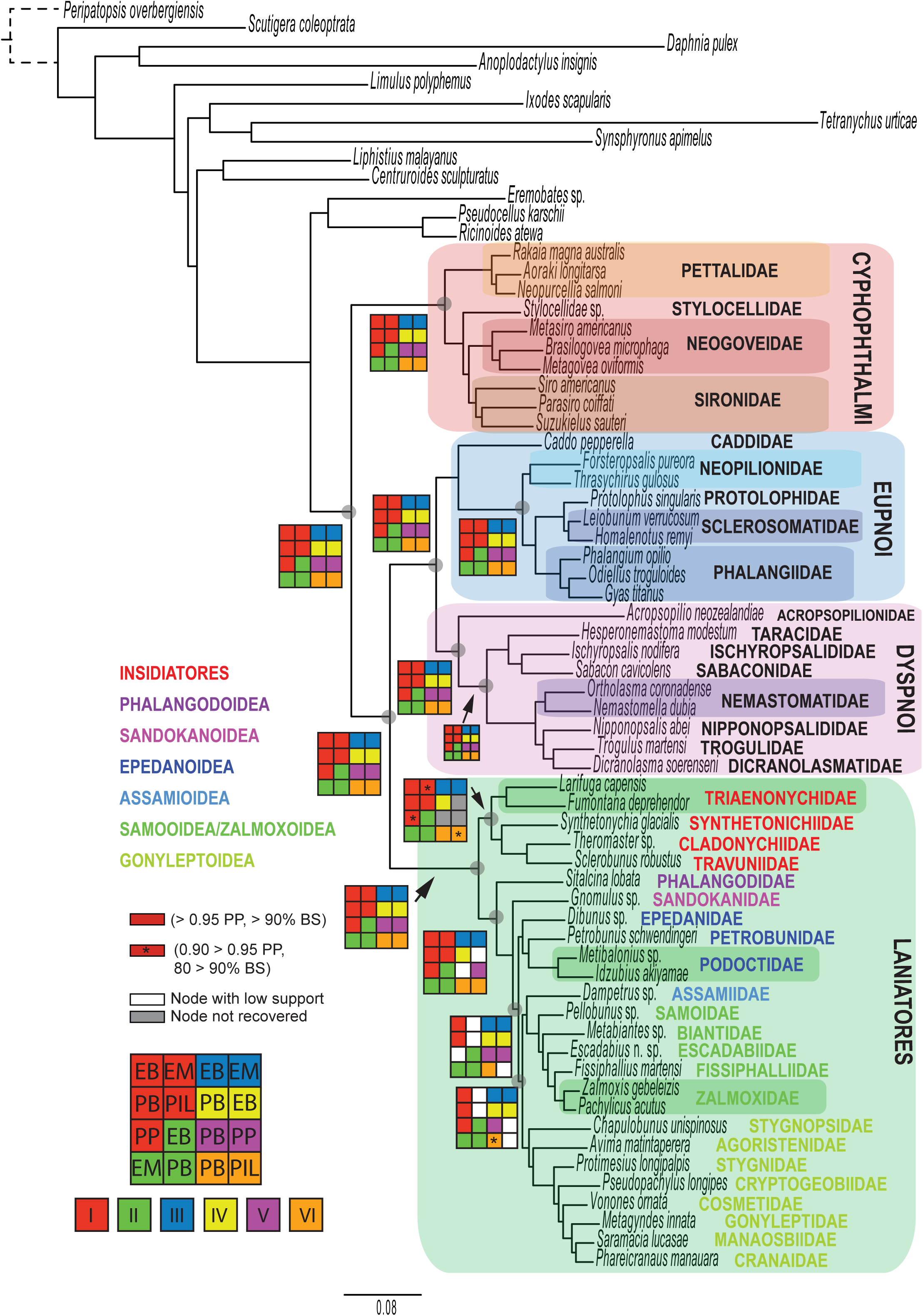
Live habitus of selected specimens: a. Synthetonychia glacialis from Westland/Tai Poutini NP, South Island, New Zealand, 19.i.2014 (MCZ 137212); b. Fumontana deprehendor from Great Smoky Mountains NP, Tennessee, USA, 1.ix.2014 (MCZ 46881); c. Gnomulus latoperculum from Gunung Ambang, Sulawesi, Indonesia, 19.vi.2006 (MCZ 131263); d. Idzubius akiyamae from Izu Peninsula, Honshu, Japan, 3.xii.2014 (MCZ 49771); e. Fissiphallius martensi from Reserva Ducke, Amazonas, Brazil, 17.xii.2013 (MCZ 136546); f. Pellobunus sp. from Isla Colón, Bocas del Toro, Panama, 22.iii.2014 (MCZ 31593); g. Avima matintaperera from Reserva Ducke, Amazonas, Brazil, 18.xii.2013 (MCZ 136560); h. Saramacia lucasae from Reserva Ducke, Amazonas, Brazil, 15.xii.2013 (MCZ 136518); i. Protimesius longipalpis from Reserva Ducke, Amazonas, Brazil, 15.xii.2013; j. Phareicranaus manauara from Reserva Ducke, Amazonas, Brazil, 17.xii.2013; k. Escadabius sp. nov. from Reserva Ducke, Amazonas, Brazil, 16.xii.2013; l. Pseudopachylus longipes from Serra de Paranapaciaba, São Paulo, Brazil, 3.vi.2014 (MCZ 32167).

The phylogeny of the order Opiliones and its four extant suborders—Cyphophthalmi, Eupnoi, Dyspnoi and Laniatores—has received considerable attention, based on morphological (Martens, 1976, 1980; Martens et al., 1981; Shultz, 1998; Giribet and Dunlop, 2005), molecular (Shultz and Regier, 2001; Giribet et al., 2010; Hedin et al., 2010; Groh and Giribet, 2015), and combined data sets (Giribet et al., 1999; Giribet et al., 2002; Garwood et al., 2014). After some debate, the relationships among the Opiliones suborders have been settled, with Cyphophthalmi constituting the sister group of Phalangida, the latter divided in Palpatores (Eupnoi + Dyspnoi) and Laniatores. More recently, a few studies have used phylogenomic data derived from transcriptomes to further test relationships among Opiliones (Hedin et al., 2012a; Sharma and Giribet, 2014; Sharma et al., 2014), but these pioneering studies included a handful of species (8 to 14) representing just a few families. Likewise, the internal relationships of each of the four suborders have received attention, mostly using molecular (Boyer et al., 2007; Sharma and Giribet, 2011; Richart and Hedin, 2013; Schönhofer et al., 2013; Groh and Giribet, 2015) and combined analyses of morphology and molecules (Giribet et al., 2012a). Other morphological analyses have focused on particular suborders (Shear, 1980; Shear and Gruber, 1983; Shear, 1986; Hunt and Cokendolpher, 1991). Recently a proposal for a Dyspnoi phylogeny, not based on data analyses, has been presented (Schönhofer, 2013). In addition, dozens of papers have explored the relationships of individual families or groups of closely related species (for some recent examples, see Schönhofer et al., 2013; Derkarabetian and Hedin, 2014; Pinto-da-Rocha et al., 2014; Vélez et al., 2014; Boyer et al., 2015; Schönhofer et al., 2015; Giribet et al., 2016; Richart et al., 2016). We can thus say that the phylogenetic understanding of Opiliones is on the right track, but some key nodes are yet to be resolved with confidence.

While many aspects of the phylogeny of Opiliones are now well understood, a few remain largely unresolved or understudied. For example, within Cyphophthalmi, the relationships among its six families, and even the monophyly of Sironidae, remain unsettled (Giribet et al., 2012a). Relationships within Eupnoi—the group that includes the true “daddy-long-legs”—are barely explored from a molecular perspective (see Hedin et al., 2012b; Vélez et al., 2014; Groh and Giribet, 2015), and no study has included all the relevant diversity. Resolution within these clades is poorly understood, with the exception of the deepest division between Caddoidea and Phalangioidea (Groh and Giribet, 2015). Relationships within Dyspnoi are beginning to settle (Richart and Hedin, 2013; Schönhofer et al., 2013; Richart et al., 2016), but for example, only recently it was recognized that Acropsopilionidae are related to Dyspnoi and not to Eupnoi (Groh and Giribet, 2015), based on a handful of Sanger-sequenced molecular markers. This resulted in transferring a clade of Opiliones from Eupnoi to Dyspnoi, as the sister group to all other members (Ischyropsalidoidea + Troguloidea), and therefore deserves further testing using a modern and more complete data set. Finally, relationships within Laniatores have changed considerably after the study of Sharma and Giribet (2011), as the taxonomy of this large clade of Opiliones has been in flux, with description of several families in recent years (Sharma and Giribet, 2011; Sharma et al., 2011; Kury, 2014; Bragagnolo et al., 2015b). Some novel results include the proposal of a sister group relationship of the New Zealand endemic family Synthetonychiidae to all other Laniatores (Giribet et al., 2010; Sharma and Giribet, 2011)—a result hinged on partial data from a single species. In addition, the relationships among many families of Insidiatores and Grassatores remain unstable.

Recent application of dense taxon sampling using large numbers of genes through modern phylogenomic approaches (e.g., based on genome and Illumina-based data sets) has resolved family-level relationships of a diversity of groups of arachnids (Bond et al., 2014; Fernández et al., 2014a; Fernández and Giribet, 2015; Sharma et al., 2015) and other arthropods (Misof et al., 2014; Fernández et al., 2016). We applied these methodologies to Opiliones phylogenetics to produce a densely sampled family-level phylogeny by analyzing 54 harvestman transcriptomes (40 newly generated for this study and 14 previously published) representing 40 of the 50 currently recognized extant families (80% familial representation).

## Materials and Methods

### Specimens

Specimens of Opiliones selected for this study were collected in a diversity of locales, including major trips to the Brazilian Amazon, Europe, North America and New Zealand, and complemented by smaller trips to other localities and by donations of specimens by colleagues. Taxonomic and geographic details of the represented species and accession codes are available in Table 1; additional information on the specimens can be found in the Museum of Comparative Zoology’s online database MCZbase (http://mczbase.mcz.harvard.edu) by using the MCZ accession numbers.

Collected specimens were preserved in RNAlater and transferred to liquid nitrogen upon arrival to the laboratory and subsequently stored at −80 ºC or, if brought alive to the laboratory, directly flash frozen in liquid nitrogen and preserved in −80 ºC. Total RNA was extracted with TRIzol (Life Sciences) and mRNA was purified with the Dynabeads mRNA Purification Kit (Invitrogen) following manufacturer’s instructions. cDNA libraries were constructed in the Apollo 324 automated system using the PrepX mRNA kit (IntegenX) and sequenced in-house at the FAS Center for Systems Biology in an Illumina Hi-Seq 2500 with a read length of 150 base pairs. Further details about the protocols can be found elsewhere (Fernández et al., 2014b; Fernández et al., 2016).

Our final matrix comprises 54 taxa, including 10 Cyphophthalmi (4 families included; Ogoveidae and Troglosironidae missing), 9 Eupnoi (representatives of all 5 families included), 9 Dyspnoi (all 8 families included), and 26 Laniatores (representatives of 23 families included; missing Gerdesiidae, Guasiniidae, Icaleptidae, Kimulidae, Metasarcidae, Nippononychidae, Pyramidopidae, and Tithaeidae, all families of relatively narrow distribution in places difficult to access). As outgroups, we included several chelicerates (*Anoplodactylus insignis, Limulus polyphemus, Synsphyronus apimelus, Tetranychus urticae, Liphistius malayanus, Eremobates sp., Centruroides sculpturatus, Ricinoides karschii* and *Pseudocellus pearsei*), one crustacean (*Daphnia pulex*), one myriapod (*Scutigera coleoptrata*) and one onychophoran (*Peripatopsis obervergiensis*) (Table 1).

All raw sequences are deposited in the SRA archive of GenBank under accession numbers specified in Table 1. Data on specimens are available from MCZbase (http://mczbase.mcz.harvard.edu).

### Orthology assignment and matrix construction

Orthology assignment was based on the OMA algorithm version 0.99.z3 (Altenhoff et al., 2013), as specified in detail in our previous work (Fernández et al., 2014b). Orthologous genes were aligned with MUSCLE (Edgar, 2004). Probabilistic alignment masking of each orthologous gene was done with ZORRO (Wu et al., 2012), and positions below a confidence threshold of 5 (see Fernández et al., 2014b) were removed from the alignments.

A first approach to matrix construction based on gene occupancy was followed (Hejnol et al., 2009). These include matrices with >90%, >75% and 50% gene occupancy (a gene occupancy of 50% means that an OMA orthologous group is selected if present in at least 50% of the taxa), resulting in three matrices with 78 genes (matrix I), 305 genes (matrix II) and 1,550 genes (matrix III), respectively.

Additionally we explored subsets of 100 genes with different evolutionary rates (measured as percentage of identical sites), as in Fernández et al. (2014a). From the 1,550-gene matrix, we selected the 100 least conserved genes (matrix IV), the 100 that had an evolutionary rate closest to the mean observed in this data set (varying from 27.9 to 28.4% of identical sites; matrix V), and the 100 least conserved genes (matrix VI). Information about the number of amino acids and the percentage of missing data for each matrix can be found in S1 Table.

### Phylogenetic analyses

Maximum likelihood inference was conducted with PhyML-PCMA (Zoller and Schneider, 2013), ExaML (Aberer and Stamatakis, 2013) and PhyML v.3.0.3 implementing the integrated branch length flag (i.e., this approach integrates branch length over a wide range of scenarios, therefore allowing implementation of a further correction of heterotachy not considered by mixture models). Bootstrap support values were estimated with 100 replicates under the rapid bootstrapping algorithm (Stamatakis et al., 2008). In PhyML-PCMA, we selected 20 principal components and empirical amino acid frequencies. The per site rate category model was selected in ExaML.

Bayesian analyses were conducted with ExaBayes (Aberer et al., 2014) and PhyloBayes MPI 1.4e (Lartillot et al., 2013) using the site-heterogeneous CAT-GTR model of evolution in the latter software (Lartillot and Philippe, 2004). Two independent Markov chain Monte Carlo (MCMC) chains were run for > 5,000 cycles. The initial 20% trees sampled in each MCMC run prior to convergence (i.e., when maximum bipartition discrepancies across chains < 0.1) were discarded as the burn-in. A 50% majority-rule consensus tree was then computed from the remaining trees sampled every 10 cycles.

Compositional homogeneity of each gene and taxon was evaluated in BaCoCa (Kück and Struck, 2014). The relative composition frequency variability (RCFV) values (that measures the absolute deviation from the mean for each amino acid for each taxon) was plotted in a heatmap using the R package gplots with an R script modified from Kück and Struck (2014). None of the genes in any of the matrices showed any signs of compositional heterogeneity (i.e., RCFV values were lower than 0.025), therefore there was no need to eliminate them from our matrices (S1 Fig).

Due to computational constraints, and in order to improve the efficiency of our analyses, not all analyses were run for all the matrices (see Figs 3–4).

**Figure 3.**
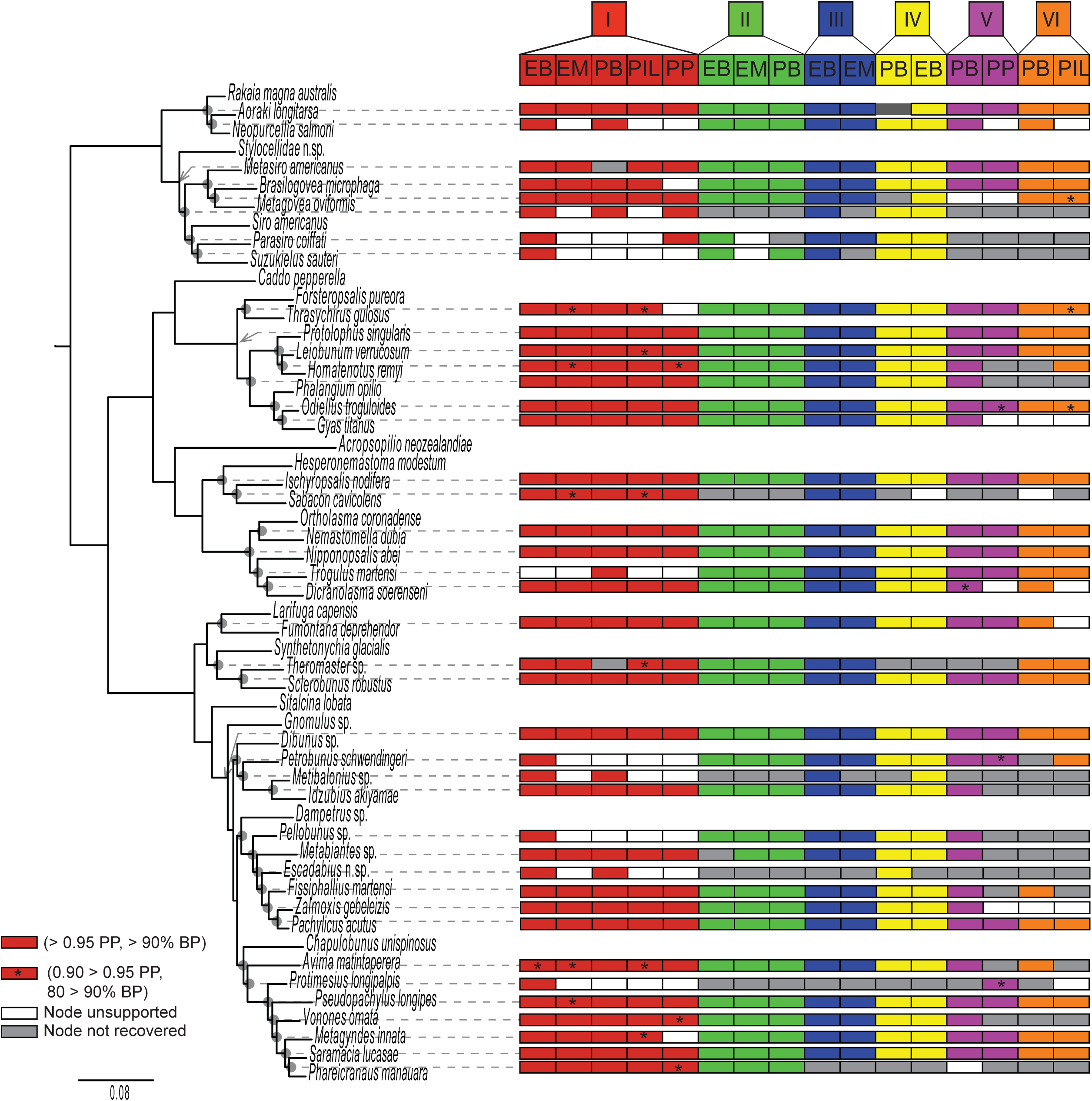
Phylogenetic hypothesis based on the 78-gene matrix I analyzed in PhyML_PCMA (-lnL = −248960.37) Selected deep nodes (gray circle) show Navajo rug illustrating support under specific data matrices and analyses. In Grassatores, colored text for family names indicate superfamily boundaries.

**Figure 4.**
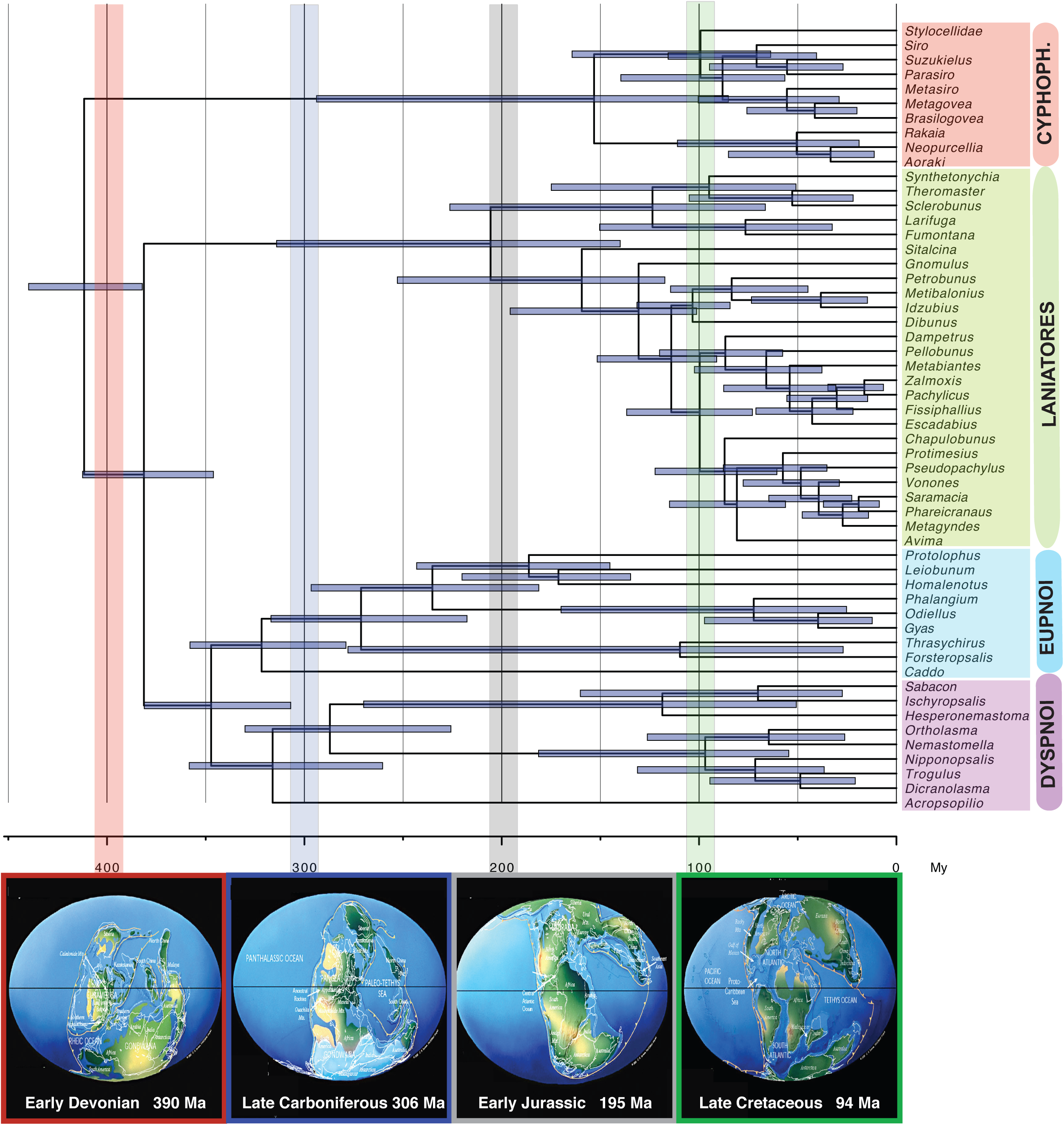
Same topology as shown in Figure 2 showcasing support for shallower nodes.

### Molecular dating

The fossil record of Opiliones is well documented, and most key fossils have been included in prior phylogenetic analyses, making their placement in a phylogenetic context precise. We mostly follow the strategy and fossil placement of Sharma and Giribet (2014), who conducted tip dating in one of their analyses. The oldest Opiliones fossil, *Eophalangium sheari*, from the Early Devonian Rhynie Cherts (Dunlop et al., 2003, 2004) is now interpreted as a member of the extant suborder Tetrophthalmi, together with *Hastocularis argus* from the Carboniferous Montceau-les-Mines Lagerstätte (Garwood et al., 2014). Tetrophthalmi is the putative sister group of Cyphophthalmi (Sharma and Giribet, 2014). *Eophalangium* had been originally interpreted as a stem Eupnoi, but now we use it as a minimum age for the origin of its sister group, Cyphophthalmi, which here corresponds to the node splitting Cyphophthalmi from Phalangida. Given how close this fossil is to the root of Opiliones we use a soft bound of 411 Ma for the floor of Opiliones. A second analysis was conducted considering the age of *Eophalangium* as the minimum age of Cyphophthalmi, based on the rationale explained above.

Two other key Carboniferous fossils (Garwood et al., 2011) are used to constrain the non-caddid Eupnoi (based on the modern-looking *Macrogyion cronus*) and the non-acropsopilionid Dyspnoi (based on *Ameticos scolos*), and applied soft bounds of 305 Ma to each of these clades.

Additional Eupnoi fossils from the Middle Jurassic (approx. 165 Mya) of Daohugou, Inner Mongolia, China are known (Huang et al., 2009), and one species, *Mesobunus dunlopi*, preserved the penis, allowing placement in Sclerosomatidae (Giribet et al., 2012b), and constraining the corresponding node to 165 Ma.

Other relevant fossils include Upper Cretaceous (lowermost Cenomanian, ca. 99 Ma) Burmese amber from Myanmar (Giribet and Dunlop, 2005; Poinar, 2008). *Petrobunoides sharmai* (Selden et al. 2016), placed in the extant family Epedanidae, was used as a constrain for the superfamily Epedanoidea. *Halitherses grimaldii* has been recently reinterpreted as a new family of uncertain affinities (Shear, 2010; Dunlop et al., 2016), and thus is of little help for dating, as an older fossil was already used to constrain the non-Acropsopilionidae Dyspnoi. *Paleosiro burmanicum* was originally described as a member of the Cyphophthalmi family Sironidae (Poinar, 2008), but it is of stylocellid affinities (Giribet et al., 2012a). Given the uncertain placement of the fossil within Stylocellidae and the presence of a single stylocellid terminal, we do not use it for node dating to avoid a “push towards the present” effect (see Giribet, 2015).

The split between Onychophora and Arthropoda was used to root the tree with a uniform prior of 528–558 Ma. The split between Xiphosura and Arachnida was constrained with a uniform prior of 465–485 Ma, based on the age of *Lunataspis aurora*. *Proscorpius osborni* was selected as being an anatomically well-understood Silurian scorpion (Whitfield 1885), setting a constraint on the split of Scorpiones from Tetrapulmonata.

Divergence dates were estimated using the Bayesian relaxed molecular clock approach as implemented in PhyloBayes v. 3.3f (Lartillot et al., 2013) under the autocorrelated lognormal and uncorrelated gamma multipliers models, resulting in four analyses (i.e., these two models were applied to both calibration configurations described above, with the age of *Eophalangium* as the minimum age of Cyphophthalmi or as the floor of Opiliones). Two independent MCMC chains were run for each analysis (10,000–12,000 cycles). The calibration constraints were used with soft bounds (Yang and Rannala, 2006) under a birth–death prior.

## Results and Discussion

All results are based on three original data matrices of 78 genes (matrix I; > 90% gene occupancy), 305 genes (matrix II; > 75% gene occupancy) and 1,550 genes (matrix III; > 50% gene occupancy), as well as subsets of these matrices (see Methods).

Figures 3–4 illustrate the topology obtained for the 78-gene matrix (matrix I) in PhyML_PCMA, with a Navajo rug representing the support for the sixteen analyses conducted for the different matrices and methods (see Materials and Methods section and figure captions).

### Higher-level Opiliones phylogenetics

Our analyses recover a stable relationship among the four extant Opiliones suborders, each well supported as monophyletic in all the analyses (Fig. 3), as consistently found in a variety of published Opiliones analyses (e.g., Shultz, 1998; Shultz and Regier, 2001; Giribet et al., 2010; Hedin et al., 2012a; Sharma and Giribet, 2014; Sharma et al., 2014; Groh and Giribet, 2015), including phylogenomic ones (Hedin et al., 2012a; Sharma and Giribet, 2014; Sharma et al., 2014). However, most published analyses found little support for the resolution within each suborder—in Sanger-based analyses due to insufficient sequence data and in phylogenomic analyses due to few taxa. The resolved relationships within each of the four suborders are thus the most novel aspects of this study. Each suborder is therefore discussed in detail below.

### Cyphophthalmi—the mite harvestmen

The members of the suborder Cyphophthalmi (Fig. 1a–b) have received especial attention phylogenetically due to their antiquity, their global distribution, and their low vagility (e.g., Boyer et al., 2007; Giribet et al., 2012a). Here we confirm the division of Cyphophthalmi into the temperate Gondwanan family Pettalidae (Fig 1a) and the remaining families (Stylocellidae, Neogoveidae, Sironidae) (Figs 3–4), a divergence that took place around the Carboniferous (Fig 5). While the New Caledonian endemic Troglosironidae (12 spp. in the genus *Troglosiro*) and the West African endemic Ogoveidae (3 spp. in the genus *Ogovea* from Cameroon, Gabon and Equatorial Guinea) were not included, their phylogenetic affinity to Neogoveidae in the clade Sternophthalmi is strongly supported by an array of morphological and molecular data sets (Giribet et al., 2012a), and is not discussed further here, as we lack phylogenomic data to test this clade.

**Figure 5.**
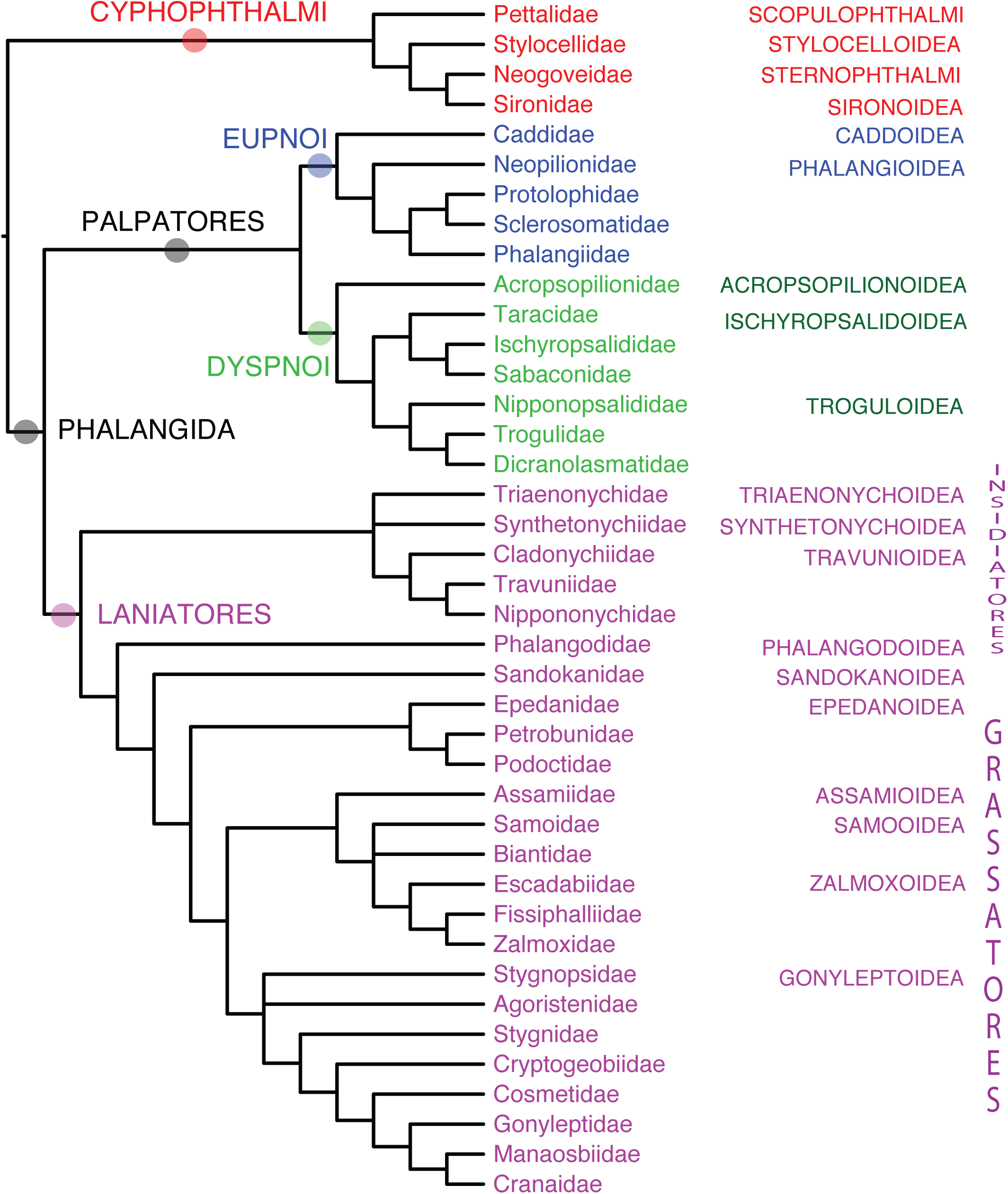
Up, chronogram of Opiliones evolution for the 78-gene data set with 95% highest posterior density (HPD) values for the dating for the first calibration configuration (i.e., the age of Eophalangium as the minimum age of Cyphophthalmi) under uncorrelated gamma model. Down, palaeogeographical reconstruction according to Christopher R. Scotese (maps modified from http://www.scotese.com/earth.htm) at some of the key ages of the split of Opiliones main lineages, as recovered by the molecular dating analysis. Vertical bars indicate correspondence with each paleomap following a color code.

Relationships among Stylocellidae, Neogoveidae and Sironidae (Fig 1b) are unstable, and two topologies prevail: (Stylocellidae, (Neogoveidae, Sironidae)) and (Sironidae, (Stylocellidae, Neogoveidae)), and neither topology supports the taxon Boreophtahlmi, grouping Stylocellidae and Sironidae, as proposed by Giribet et al. (2012a) (Figs 3–4). The first topology is preferred by the most-complete data set. However, in some of the analyses using fewer genes the Stylocellidae species nest within Sironidae, albeit without support. These two alternatives will require further examination with more stylocellid samples, as the alternative topologies may also have an impact on the dating, which suggests an initial diversification around the Jurassic/Cretaceous boundary (Fig. 5).

Monophyly of Neogoveidae is recovered in all analyses, with the exception of the PhyloBayes analysis of the 78-gene matrix (Fig. 5). The placement of the North American *Metasiro* with the typical Neotropical neogoveids corroborates previous molecular hypotheses of the delimitation of this clade (Giribet et al., 2012a; Benavides and Giribet, 2013).

Monophyly of Sironidae (Fig. 1b), here represented by three genera of the three main lineages of this Laurasian family (*Siro, Parasiro*, and *Suzukielus*) is unstable across analyses (Fig. 4). Monophyly of Sironidae has been difficult to obtain in molecular analyses as well as with morphology due to differences in family-level characters in the Western Mediterranean *Parasiro* and the Japanese *Suzukielus* (Giribet et al., 2012a).

### Eupnoi—the daddy-long-legs

Family-level Eupnoi phylogenies are scarce (Giribet et al., 2012a; Hedin et al., 2012b; Groh and Giribet, 2015) and have typically undersampled southern hemisphere lineages. Our analyses support the well-known division of Caddoidea (Fig. 1c) and Phalangioidea (Fig. 1d–f), which in turn divides into the southern hemisphere Neopilionidae (Fig. 1d) and the mostly northern hemisphere families Phalangiidae (Fig. 1e), Sclerosomatidae (Fig. 1f) and Protolophidae—although Phalangiidae and Sclerosomatidae have later diversified in the Southern hemisphere (Figs 3–4). The sister group relationship among Protolophidae and Sclerosomatidae has been found in previous analyses (Hedin et al., 2012b; Groh and Giribet, 2015), and in fact some have considered Protolophidae a junior synonym of Sclerosomatidae (Kury, 2013). However, resolution among the families of Phalangioidea has received little or no support in previous studies. Our results thus provide, for the first time, a well-resolved Eupnoi phylogeny, including the placement of *Gyas titanus* within Phalangiidae, as suggested by Hedin et al. (2012b), instead of Sclerosomatidae. Unfortunately we were not able to include any members of the phylogenetically unstable "*Metopilio* group" (Giribet et al., 2010; Hedin et al., 2012b). We thus find Phalangioidea divided into three main clades: Neopilionidae, Sclerosomatidae/Protolophidae, and Phalangiidae (including *Gyas*). However, the systematics of this large group of Opiliones, with nearly 200 genera and 1,800 species, will require much denser sampling before the group can be properly revised.

The division between the temperate Gondwanan Neopilionidae and the remaining species is well supported phylogenetically and biogeographically, as it is inferred to be Permian (Fig. 5).

### Dyspnoi

The global phylogeny of Dyspnoi has received attention from different workers using morphology and molecules, but only recently there has been modern treatment. Groh and Giribet (2015) finally circumscribed the suborder, transferring Acropsopilionidae (Fig. 1g) from Eupnoi to the sister group of all other Dyspnoi (Ischyropsalidoidea [Fig. 1h] and Troguloidea [Fig. 1i]) based on molecular data analyses of a few Sanger-sequencing genes and morphological examination. Our phylogenomic datasets corroborate this topology, placing *Acropsopilio neozealandiae* (Fig. 1g) as the sister group to the other Dyspnoi, with the monophyly of each of Ischyropsalidoidea and Troguloidea being fully supported. While the position of *Halitherses grimaldii* remains uncertain, their large eyes, resembling those of caddids and acropsopilionids, and their troguloid facies (Shear, 2010) suggest a phylogenetic placement between Acropsopilionoidea and the remaining Dyspnoi (Dunlop et al., 2016), perhaps as sister group to Troguloidea or to Troguloidea + Ischyropsalidoidea, although the caddoid eyes are now found in Caddoidea, Phalangioidea (in the members of the genus *Hesperonemastoma*; see (Groh and Giribet, 2015)), and Acropsopilionoidea, and are thus best optimized as a symplesiomorphy of Palpatores.

The main division in Dyspnoi follows a biogeographic pattern, with Acropsopilionoidea mostly concentrating in the southern hemisphere, while the remaining Dyspnoi being circumscribed to Laurasia, a biogeographic pattern consistent the Carboniferous/Devonian split between these two lineages (Fig. 5).

### Laniatores—the armored harvestmen

The phylogeny of Laniatores has received recent attention at many levels (Giribet et al., 2010; Sharma and Giribet, 2011; Cruz-López et al., 2016; Sharma et al., 2016). In an unpublished thesis, Kury (1993) divided Laniatores into Insidiatores (Fig 2a–b) and Grassatores (Fig 2c–l), a division found here, but not in some studies that included a meaningful sampling of Laniatores (Giribet et al., 2010; Sharma and Giribet, 2011). These instead found the New Zealand endemic Synthetonychiidae (Fig 2a) to be the sister group to all other Laniatores (Eulaniatores *sensu* Kury (2015)), a result worth testing due to the partial nature of the included synthetonychiid sequences and the former placement of this family as sister group to Triaenonychidae. Of special interest was also the phylogenetic position of the unstable North American *Fumontana deprehendor* (Fig. 2b), a member of the mostly temperate Gondwanan family Triaenonychidae that was poorly resolved in prior studies. Our analyses do find a sister group of *Fumontana* to the representative of the southern hemisphere Triaenonychidae in virtually all analyses, with a Permian divergence (Fig. 5). *Synthetonychia* is either the sister group of the represented travunioids in most analyses (except for matrices IV and V; see Fig. 4), or sister group to Triaenonychidae in some analyses, as originally proposed by Forster (1954). Further discussion on Insidiatores will require increased diversity of genera both within Triaenonychidae and within the travunioid families (see for example Kury, 2013).

Resolution within Grassatores has remained elusive except for the recognition of a main division between Phalangodidae and the remaining Grassatores and of the superfamilies Gonyleptoidea, Assamioidea, Zalmoxoidea and Samooidea, and perhaps a clade of Southeast Asian families, Epedanoidea (Sharma and Giribet, 2011), although some of these clades were not supported in re-analyses of the Sharma & Giribet dataset (Cruz-López et al., 2016; Sharma et al., 2016). Here we find Phalangodidae (represented by *Sitalcina lobata*) to be the sister group of the remaining Grassatores, a result supported in all analyses conducted to date.

The SE Asian endemic Sandokanidae (Fig 2c) (Sharma and Giribet, 2009) is resolved as the sister group to the remaining families, the latter clade being supported by nearly all matrices and most analyses (only some analyses find this clade without support) (Fig 3). The position of Sandokanidae with respect to the other Grassatores families has been difficult to resolve in prior analyses (Giribet et al., 2010; Sharma and Giribet, 2011), which sometimes suggested a relationship to Epedanoidea. Here we reject this hypothesis and support Sandokanidae as the second offshoot of the Grassatores clade, i.e., (Phalangodidae, (Sandokanidae, remaining Grassatores)).

The sister group of Sandokanidae divide into the largely SE Asian Epedanoidea (represented here by members of Epedanidae, Petrobunidae and Podoctidae) and a clade including Assamioidea, Samooidea, Zalmoxoidea and Gonyleptoidea, this divergence being Cretaceous (Figs 6–7). This coincides with the first Laniatores fossils, which were already present in the terranes of today’s Myanmar (Selden et al., 2016). Epedanoidea is monophyletic in all analyses (although sometimes without significant support), except for the PhyloBayes analysis of matrix VI (Fig. 4), and it is resolved with Epedanidae being sister group to a clade of Petrobunidae and Podoctidae (Fig 2d), although support for these relationships is not optimal (Fig 4; see also Sharma et al., 2016), as it may require additional taxa, including the missing family Tithaeidae and additional species. Epedanoidea has however been difficult to recover in a recent analysis focusing on Podoctidae (Sharma et al., 2016).

**Figure 6.**
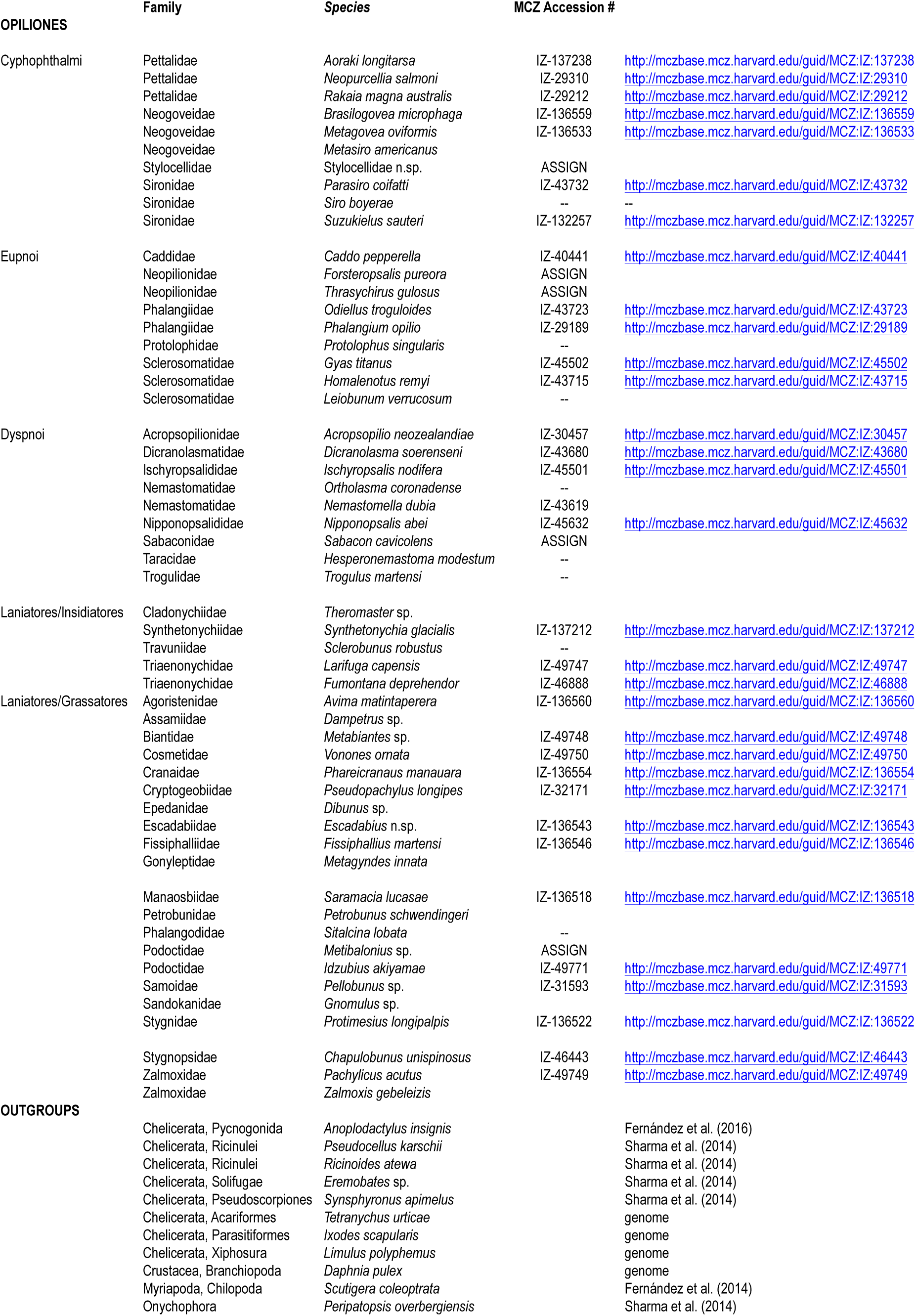
Summary tree of familial and superfamilial relationships of Opiliones supported in this study, with major nodes highlighted.

The sister group of Epedanoidea, a clade composed of Assamioidea–Zalmoxoidea– Samooidea–Gonyleptoidea is well supported in virtually all analyses (Fig 3). Internal resolution among these superfamilies had found conflict in prior studies (Giribet et al., 2010; Sharma and Giribet, 2011), as it probably required additional data to resolve this rapid radiation of Laniatores families. Phylogenomic data find the much-needed information to resolve this clade, here assigned a Cretaceous age (Figs 6-7).

A sister group relationship of *Dampetrus* (an assamiid, the only representative of Assamioidea included here) to Zalmoxoidea-Samooidea is found with all data matrices except V and VI, but does not receive support in most analyses with matrix I. Fewer genes seem to be necessary to support fully a clade of Zalmoxoidea and Samooidea, which was found in prior Sanger-based studies (Sharma and Giribet, 2011). However, Samooidea is paraphyletic with respect to Zalmoxoidea in about half of the analyses (Fig. 4), although in others both superfamilies are reciprocally monophyletic. The *bona fide* samooid *Pellobunus* from Panama (Fig 2f) is sister group to the remaining members of this clade, followed by a representative of biantidae *Metabiantes* from South Africa, and by a clade of Zalmoxoidea, including an Amazonian specimen we tentatively placed in the genus *Escadabius* (Escadabiidae) (Fig 2k), and then the representatives of Fissiphalliidae (*Fissiphallius martensi*; Fig 2e) and Zalmoxidae (*Zalmoxis, Pachylicus*). Except for the clade placing Metabiantes with the zalmoxoids, relationships within this clade are stable (Fig 4). Further samooid and zalmoxoid missing families (Kimulidae, Stygnommatidae, Guasiniidae and Icaleptidae; all exclusively Neotropical) should be sampled to resolve the issue of the reciprocal monophyly of the families, and in fact, when a second stygnopsid was evaluated in the expanded data set (*Karos*), Samooidea was supported as a clade (Fig 5).

Gonyleptoidea (Fig 2g—j) is restricted to an expanded Neotropics (some species of gonyleptid make it into Patagonia and some cosmetids quite North into the USA). Stygnopsidae (*Chapulobunus*) is well supported as sister group to all other gonyleptoids, followed by Agoristenidae (*Avima*; Fig 2g), although in this case the position of Agoristenidae is not well resolved. Agoristenids had been proposed as the sister group of the non-stygnopsid gonyleptoids in previous analyses (Sharma and Giribet, 2011), as suggested here in some trees, but most matrices suggest a sister group relationship of Agoristenidae and Stygnopsidae. Stygnidae (*Protimesius*; Fig 2i) is well supported as sister group to all the remaining families, followed in a ladder-like fashion by the families Cryptogeobiidae (*Pseudopachylus*; Fig 2l), Cosmetidae (*Vonones*), Gonyleptidae (*Metagyndes*), Manaosbiidae (*Saramacia*; Fig 2h) and Cranaidae (*Phareicranaus*; Fig 2j). This topology is fully compatible with more detailed recent analyses of Gonyleptoidea (Kury, 2014; Pinto-da-Rocha et al., 2014; Bragagnolo et al., 2015a), some of which consider Manaosbiidae and Cranaidae junior synonyms of Gonyleptidae (Pinto-da-Rocha et al., 2014), although this synonymy was not accepted in subsequent studies (Kury, 2014; Kury and Villarreal M, 2015). We thus support a clade including these three families, which has also been called GG or Greater Gonyleptidae (Kury, 2014; Kury and Villarreal M, 2015), but maintain the three families until this issue gets resolved.

Grassatores are much more abundant in the southern than in the northern hemisphere, and thus some authors have seen the former as a center of origin of the group. However, our analyses consistently resolve northern hemisphere clades at the base of the Grasssatores tree (Figs 6-7), with the widespread family Phalangodidae splitting first, followed by the SE Asian Sandokanidae. The next clade in the tree divides into Epedanoidea, comprising mostly SE Asian and Philippine families and a largely Neotropical clade (from the southern part of Gondwana). The southern hemisphere clade has extended into the northern hemisphere in a few cases, especially Cosmetidae, Podoctidae, Samoidae and Zalmoxidae, the latter three families well known for the ability of dispersing across oceanic barriers (Sharma and Giribet, 2012). While denser sampling should test the origin of Cosmetidae and Samoidae, it is clear that Zalmoxidae originated in the Neotropics.

### Molecular dating and general biogeographical patterns in Opiliones

The molecular dating analyses for both calibration configurations (i.e., the age of *Eophalangium* as the floor of Opiliones or Cyphophthalmi) yielded very similar results, with the main differences obtained between the autocorrelated and uncorrelated model analyses within each calibration configuration (Fig S2). For purposes of conservatism, we discuss our results based on the chronogram under the uncorrelated gamma multipliers model using the age of *Eophalangium* as the minimum age of Cyphophthalmi (Fig 5).

Opiliones have been often used as examples of animals with ancient and conservative biogeographic patterns, therefore suitable for vicariance biogeographic analyses (Giribet and Kury, 2007; Giribet et al., 2010). One general pattern observed here is a division between temperate Gondwana (the terranes that were once directly connected to Antarctica) and the remaining landmasses, including, in some cases, clades currently in tropical Gondwana. This is for example the case of Cyphophthalmi, with a main division between the strictly temperate Gondwanan family Pettalidae and the remaining families (this including Laurasian and tropical Gondwanan clades), or in Dyspnoi, with Acropsopilionidae, being mostly distributed in temperate Gondwana, as the sister group to the rest of the families, restricted to the northern hemisphere. Within Eupnoi Caddidae is mostly Laurasian, but Phalangida once more divides into Neopilionidae, restricted to temperate Gondwana, and the remaining families, mostly Laurasian, although some secondarily extending southwards. Once more, Insidiatores, although somehow unresolved, finds a division between the predominantly temperate Gondwanan family Triaenonychidae (or Triaenonychidae + the New Zealand endemic Synthetonychiidae) and the northern hemisphere Insidiatores (Travunioidea). In addition, Triaenonychidae has a basal split between the northern hemisphere *Fumontana* and the temperate Gondwanan clade (although here it is represented by a single species). Further biogeographic studies on Triaenonychidae are currently underway. Laniatores depict several other interesting patterns, including two clades of southeast Asian families, including Sandokanidae and Epedanoidea while the remaining species mostly appear to be of Tropical Gondwanan origins, with some remarkable cases of range expansions (e.g., trans-continental disjunctions in Assamiidae, Biantidae, Pyramidopidae, and Zalmoxidae; Sharma and Giribet, 2011, 2012; Cruz-López et al., 2016).

Interestingly, the splits between Temperate Gondwana and the rest precede the breakup of Pangea (Fig 5), suggesting ancient regionalization across Pangea, as shown in other groups of terrestrial invertebrates (Murienne et al., 2014) and in the early diversification of amphibians (San Mauro et al., 2005). Splits between tropical Gondwana and Laurasia, both in Cyphophthalmi and in Grassatores, seem to be much younger, and may be associated with the breakup of Pangea, possibly representing true Gondwanan/Laurasian vicariant events, and not the result of ancient cladogenesis and Pangean regionalization. This pattern would be more in agreement with the diversification of larger and more vagile organisms (e.g., dytiscid beetles; Bukontaite et al., 2014).

### Conclusions

Our analysis of a large number of novel transcriptomes has allowed us to propose a stable phylogeny of Opiliones (Fig 6). Such analyses of large data matrices have allowed us to place all superfamilies of Opiliones (and 80% of the families) in a resolved phylogenetic context, with only a few spots to be sorted out, these in areas of the tree of life where sampling was still limited. Our trees support most traditional relationships within Opiliones and resolve some recalcitrant familial relationships, such as a well-resolved Eupnoi phylogeny, the rejection of Boreophthalmi, the monophyly of Insidiatores, the monophyly of Triaenonychidae, and the placement of Stygnopsidae as the most basal family of Gonyleptoidea, among others. We also show that Opiliones exhibit some splits reflecting ancestral Pangean regionalization, whereas others conform with high fidelity to the sequence of Pangean fragmentation, therefore constituting ideal model systems to understand ancient biogeographic patterns.

## Acknowledgements

Many colleagues assisted with field work and samples, and we are indebted to all of them for their contribution of specimens and expertise, especially Jimmy Cabra, Ron Clouse, Pío Colmenares, Jesús Alberto Cruz, Óscar Francke, Guilherme Gainett, Abel Pérez González, Gustavo Hormiga, Carlos Prieto, Ricardo Pinto-da-Rocha, Cristiano Sampaio Porto, Willians Porto, and Nobuo Tsurusaki. The following funding sources were used: NSF grant DEB-1457539: Collaborative Proposal: Phylogeny and diversification of the orb weaving spiders (Orbiculariae, Araneae) (GG); National Geographic Society Grant #9043-11: Re-descovering the mite harvestmen (Arachnida, Opiliones, Cyphophthalmi, Neogoveidae) of the Brazilian Amazonas (GG). Fieldwork to Chile and New Zealand was supported by Putnam expedition grants from the MCZ (GG RF); fieldwork to Reserva Ducke was supported by CAPES (ALT GG); fieldwork in the Philippines and Australia was supported by NSF DBI-1202751 to PPS, and in the Philippines by a National Geographic exploration grant to Ronald M. Clouse and PPS.

## Author contributions

Collected samples: RF, PPS, ALT, GG. Performed laboratory work and analyses: RF. Wrote the paper: RF, PPS, GG.

## Supporting Information

**Figure S1.** BaCoCa results showing compositional homogeneity values per taxon and per gene in supermatrices I (page 1), IV (page 2), V (page 3), VI (page 4) and II (page %)(results in supermatrix III not shown due to a lack of resolution related to the high number of genes included in the matrix).

**Figure S2.** Chronogram of Opiliones evolution for the 78-gene data set with 95% highest posterior density (HPD) values for a) the dating for the first calibration configuration (i.e., the age of Eophalangium as the minimum age of Cyphophthalmi) under the autocorrelated (page 1) and uncorrelated gamma model (page 2), and b) the second calibration configuration (i.e., the age of Eophalangium as the floor of Opiliones) under the autocorrelated (page3) and uncorrelated gamma model (page 4).

